# DOMINO: Learning Domain Co-occurrence for Multidomain Protein Design

**DOI:** 10.64898/2026.05.01.721929

**Authors:** Fengyuan Dai, Jin Su, Qingsong Tan, Huijiao Yang, Xibin Zhou, Fajie Yuan

## Abstract

Multidomain proteins arise through the reuse and recombination of structural domains, yet natural architectures represent a sparse, structured sample of the possible domain-combination space. Here, we introduce DOMINO, a two-stage framework that learns domain co-occurrence from TED-annotated multidomain proteins and uses the learned patterns to generate new multidomain sequences. DOMIN, a contrastive retrieval model, embeds domains into a latent compatibility space and retrieves candidate partners for a query domain from a TED-derived domain pool, including pairings not observed in the TED-derived co-occurrence set. DOMO, a conditional autoregressive sequence model, converts each retrieved domain pair into a full-length protein sequence by jointly generating the specified domain regions and the non-domain sequence context between and around them. DOMIN recovers hierarchical patterns of natural domain co-occurrence and expands the observed CATH homologous-superfamily co-occurrence network with candidate novel pairings. DOMO realizes both held-out natural pairs and DOMIN-retrieved pairs as proteins with high domain recovery and high AlphaFold-predicted structural confidence. Applied at scale, DOMINO generated 5 million retrieval-derived multidomain proteins, with sampled designs showing recovery of the specified domains, diverse CATH annotations, and sequence novelty relative to UniRef100. Together, these results support domain co-occurrence as a predictive design prior and demonstrate a scalable strategy for exploring multidomain protein architectures through new combinations of existing structural modules.

## 1 Introduction

Protein evolution samples sequence space through genetic changes that span multiple scales. Point mutations act at the residue level to tune protein properties such as activity, specificity and stability [1–3], whereas larger events— including duplication, insertion, deletion, fusion, fission and recombination—reorganize longer sequence segments and diversify protein architectures [4]. These architectural changes are often naturally described in terms of protein domains: compact structural units that frequently serve as functional and evolutionary modules and can fold semi-independently [5–7]. Domains thus provide a modular framework for protein evolution, allowing existing structural units to be reused, reordered, duplicated or combined with new partners to produce new regulatory, spatial and functional properties.

Multidomain proteins illustrate that domain modularity is constrained rather than freely combinatorial. Although fusion, recombination and related genetic mechanisms can place diverse domains within the same polypeptide chain, only a small subset of possible combinations is observed and retained in extant proteins. Viable architectures must satisfy biophysical constraints imposed by domain folding, chain connectivity, linker geometry and inter-domain contacts. Evolutionarily recurrent architectures are often additionally supported by functional coupling through shared localization, pathway membership, regulation or direct physical contact [7–10]. Such recurrent combinations can themselves behave as evolutionary units larger than individual domains, forming “supradomains” that reflect structural and functional compatibility across protein families [11]. Thus, natural domain co-occurrence contains statistical signals not only of historical domain combinations, but also of the constraints that make particular recombinations viable.

These observations raise a broader question: can the rules implicit in natural domain co-occurrence be learned in a predictive form? Previous studies have systematically catalogued protein domain architectures [12] and modeled domain co-occurrence relationships as graphs [13], revealing non-random and constrained patterns in how domains are combined across protein families [14]. However, these patterns have predominantly been used for descriptive analyses of natural evolution, with limited efforts to develop predictive models of domain-pair plausibility or co-occurrence propensity. A predictive representation of domain co-occurrence could help identify likely partners for a given domain, distinguish recurrent or plausible combinations from arbitrary pairings, and generate hypotheses about how structural, functional, and evolutionary constraints shape multidomain protein organization. Such a representation could also provide a principled basis for designing new multidomain architectures, but this requires connecting domain-level pairing predictions to sequence-level protein realization.

This question has become newly accessible because protein structures and domain annotations are now available at unprecedented scale. Structural classification resources such as SCOPe [15] and CATH [6] have long organized protein domains according to structural and evolutionary relationships, but their coverage was historically limited by the number of experimentally solved structures in the Protein Data Bank [16]. Recent structure prediction methods, including AlphaFold 2/3 [17, 18], together with the AlphaFold Protein Structure Database [19], have greatly expanded structural coverage across protein sequence space. The Encyclopedia of Domains (TED) [20] further decomposes this predicted structural universe into domain-level annotations across hundreds of millions of proteins, revealing extensive domain diversity, multidomain organization and previously unobserved domain-superfamily interactions. These resources transform domain organization from a sparse catalogue into a large-scale statistical system, making it possible to learn domain co-occurrence as a source of evolutionary, structural and functional constraints.

Despite this progress, it remains unclear whether natural domain co-occurrence can be represented in a form that supports both partner inference and the construction of full-length multidomain proteins. Although deep learning has advanced protein structure prediction and design [21–25], most approaches operate primarily at the level of sequences, structures or local functional motifs, rather than explicitly learning domain-level compatibility across natural multidomain architectures. Here, we introduce DOMINO (DOMain co-occurrence INference and multidomain prOtein generation), a two-stage framework that links domain-level compatibility learning with sequence-level generation. First, DOMIN learns a latent compatibility space from natural multidomain proteins, enabling retrieval of candidate domain partners for a query domain, including combinations not observed during training. Second, DOMO uses the inferred domain specifications as conditioning information to generate full-length protein sequences, transforming proposed domain combinations into coherent multidomain chains. We show that DOMIN recovers hierarchical patterns of natural domain co-occurrence and identifies structurally plausible domain partners, and that DOMO realizes both natural and retrieval-derived specifications as foldable, diverse and sequence-novel multidomain designs. Together, DOMINO provides a framework for learning the organization of multidomain proteins and using the learned compatibility patterns to guide synthetic sequence generation.

## 2 Architecture

DOMINO separates multidomain protein design into two coupled inference problems: selecting compatible domain partners and realizing the resulting specification as a single polypeptide sequence (Fig. 1). Given a query domain, DOMIN retrieves candidate partners from a TED-derived domain pool. The selected query-target pair is then passed to DOMO, which generates a full-length multidomain sequence conditioned on the two domain specifications.

**Fig. 1.**
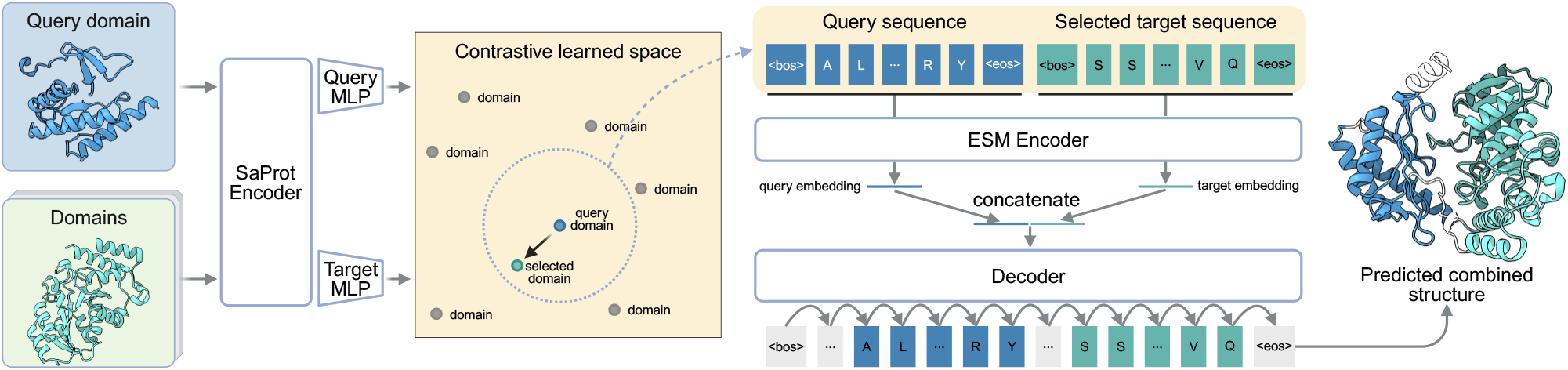
Overview of the DOMINO framework. A query domain and candidate TED domains are encoded by SaProt and projected into a contrastively learned compatibility space by DOMIN. Retrieved domain partners are paired with the query domain and supplied to DOMO as residue-level conditioning information. DOMO encodes the query and selected target sequences with an ESM encoder and uses an autoregressive decoder to generate a full-length multidomain protein sequence, including domain regions and intervening sequence context.

DOMIN is a contrastive retrieval model over protein domains. Query and candidate domains are encoded by a shared SaProt [26] encoder and mapped through separate query and target projection heads into a common compatibility space. SaProt is used in this stage because DOMIN performs retrieval over TED-derived domains with predefined boundaries and available structural context. In this setting, structure-aware representations can capture fold-level features that complement sequence patterns and are useful for compatibility retrieval. Training pairs are derived from TED-annotated natural multidomain proteins, with domains from the same protein treated as positives and non-co-occurring domains used as negatives. The contrastive objective organizes the latent space according to natural domain co-occurrence rather than sequence identity alone. At inference, DOMIN ranks candidate TED domains by similarity to the query embedding in this learned space, producing putative compatible partners for downstream generation.

DOMO is a conditional autoregressive sequence model that realizes a selected domain pair as a full-length protein. In contrast to DOMIN, DOMO uses a sequence-only ESM [27] encoder to construct residue-level conditioning embeddings for the query and target domain sequences. This choice reflects the different nature of the generation problem: the model must generate not only the specified domain regions, but also linkers and other intervening sequence context whose structures are not known before generation and may depend on the final combined chain. Sequence-based conditioning therefore provides residue-level information from the selected domains without requiring structural annotations for regions that are generated de novo. The query and target domain sequences are separately tokenized and encoded with the ESM encoder. Their residue-level embeddings are concatenated to form a joint domain specification, which is supplied to an autoregressive decoder that generates the output sequence token by token. Because the decoder attends to domain-level conditioning rather than copying a preassembled concatenation, DOMO can generate the domain regions together with linkers and sequence context required for a coherent multidomain chain.

Together, these modules define a hierarchical design procedure in which domain architecture is inferred before sequence realization. DOMIN narrows the combinatorial space of possible multidomain architectures using a learned co-occurrence prior, and DOMO translates each selected architecture into a complete sequence without relying on a fixed concatenation template. This separation creates a controllable interface between domain-level organization and sequence-level generation, enabling multidomain proteins to be designed from modular domain specifications.

## 3 Retrieving and expanding domain co-occurrence patterns with DOMIN

We evaluated DOMIN as a domain-partner retrieval model on a held-out test set using an all-to-all retrieval protocol (see Section A.1 for dataset construction details). For each query domain, DOMIN ranked all remaining test-set domains according to their predicted co-occurrence compatibility. The annotated partner domains from the same multidomain protein were treated as ground-truth partners. Unless otherwise stated, retrieval correctness was defined at the CATH homologous-superfamily level: a retrieved domain was counted as a correct hit only if its four-level CATH classification (C.A.T.H) matched that of an annotated ground-truth partner domain. This criterion evaluates whether DOMIN recovers the correct partner domain type rather than the identical domain instance.

We first analyzed top-1 retrieval results across the CATH hierarchy as a measure of structural relatedness between retrieved and annotated partner domains. As shown in Fig. 2a, 38.4% of top-1 retrieved domains matched the ground-truth partner at the Homology level (C.A.T.H). Match rates increased at progressively coarser levels of the hierarchy, reaching 59.5% at the Topology level (C.A.T), 68.7% at the Architecture level (C.A), and 77.8% at the Class level (C). These coarse-level matches should not be interpreted as retrieval accuracy in the strict sense, but rather as evidence that DOMIN tends to retrieve domains with related structural classification even when the exact homologous superfamily is not recovered.

**Fig. 2.**
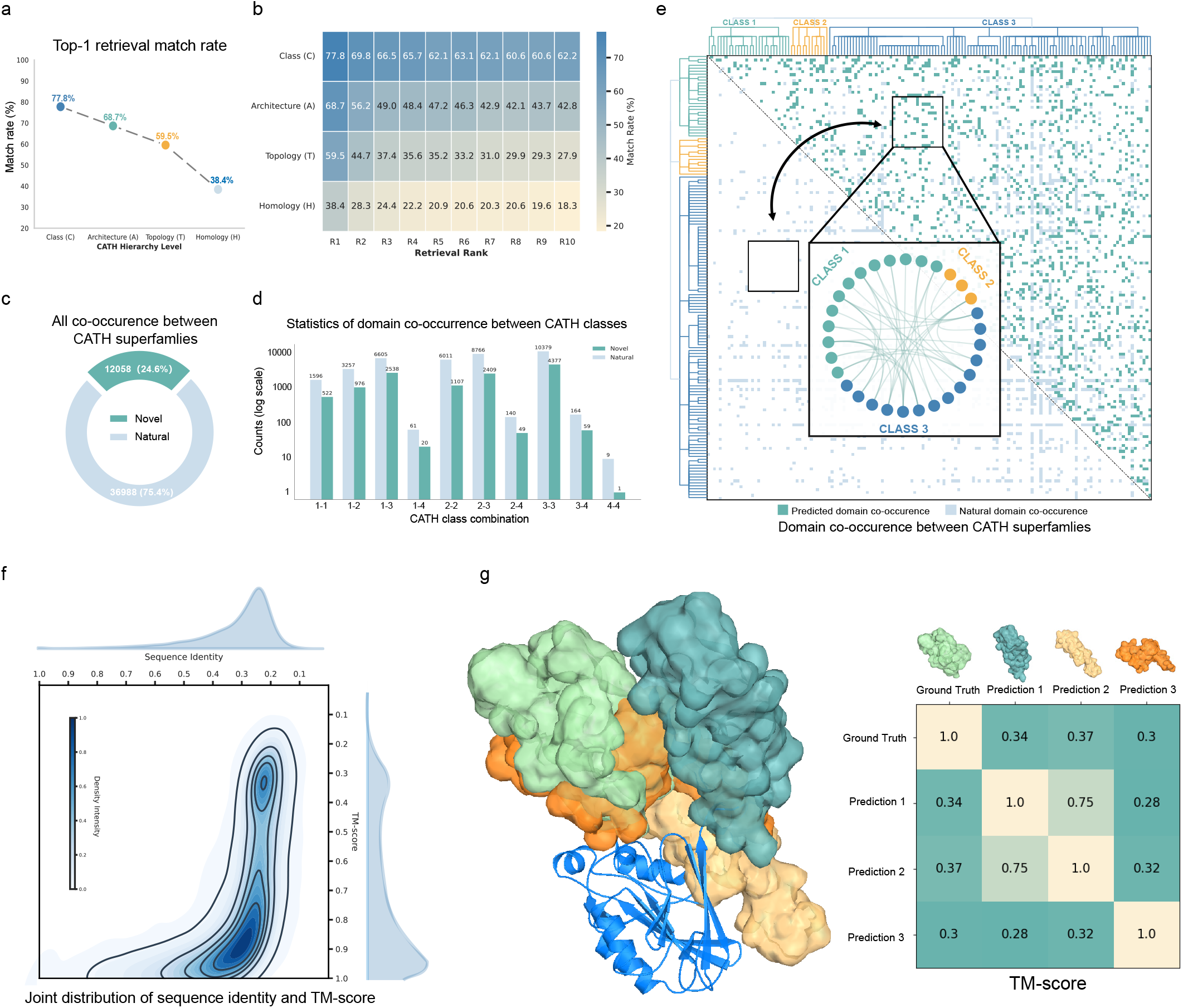
Retrieval performance and co-occurrence analysis of DOMIN. **a**, Top-1 hierarchical match rate between retrieved domains and TED-annotated ground-truth partner domains across CATH classification levels. A match at the Homology level corresponds to identical four-level CATH classification (C.A.T.H), whereas matches at Class, Architecture, and Topology indicate progressively coarser structural agreement. **b**, Rank-wise hierarchical match rate from rank 1 to rank 10 for each CATH level. Each cell reports the fraction of queries for which the domain retrieved at the corresponding rank matches the annotated partner at the indicated CATH level. **c**, Composition of CATH homologous-superfamily co-occurrence pairs, where “Novel” denotes DOMIN-predicted candidate novel co-occurrences not observed in the TED-derived natural co-occurrence set. **d**, Distribution of natural and DOMIN-predicted candidate novel co-occurrences across CATH class combinations; for example, 1–2 denotes co-occurrence between domains from CATH Class 1 and Class 2. **e**, Adjacency-matrix representation of CATH homologous-superfamily co-occurrences. Light blue denotes TED-derived natural co-occurrences, and teal denotes DOMIN-predicted candidate novel co-occurrences not observed in the TED-derived natural set. The inset highlights local connectivity patterns. **f**, Joint distribution of sequence identity versus TM-score, comparing TED-observed domains with DOMIN predictions for each query domain. **g**, Case study illustrating the query domain (blue backbone) together with paired domains (ground truth and model prediction) in distinct colors. Average pLDDT values for the full-length proteins (incorporating the query domain) are 88.1, 75.5, 76.3, and 82.7 for Ground Truth and Predictions 1–3, respectively.

To examine how retrieval quality changes across the ranked list, we computed rank-wise hierarchical match rates for the top ten retrieved domains (Fig. 2b). The rank-1 Homology-level match rate was 38.4%, corresponding to the strict C.A.T.H retrieval criterion. Homology-level match rates generally declined with retrieval rank, decreasing from 38.4% at R1 to 18.3% at R10. However, matches at broader CATH levels remained common across the top ten candidates, particularly at the Class and Architecture levels. Thus, even when a retrieved domain did not match the annotated partner at the Homology level, it often remained structurally related to the annotated partner at broader levels of the CATH hierarchy. These results indicate that DOMIN enriches the retrieved set for domains structurally related to the annotated partner, including Homology-level matches in a substantial fraction of top-ranked predictions. Because CATH-H superfamilies can contain functionally diverse members and because many biologically plausible partners may be absent from the annotated natural architecture, these metrics should be interpreted as conservative measures of superfamily-level co-occurrence recovery.

We next analyzed domain co-occurrence patterns at the level of CATH homologous superfamilies. Natural co-occurrences were defined as pairs of CATH homologous superfamilies observed within the same multidomain protein architecture based on TED domain segmentation [20]. To identify DOMIN-predicted candidate novel co-occurrences, we performed all-to-all retrieval on the retrieval database (see Section A.2), using each domain as a query and recording its top-1 retrieved domain. Each query-retrieval pair was mapped to the corresponding pair of CATH homologous superfamilies and compared with the TED-derived natural co-occurrence set. As shown in Fig. 2c, the combined co-occurrence set contained 36,988 TED-derived natural pairs (75.4%) and 12,058 DOMIN-predicted candidate novel pairs not observed in the TED-derived set (24.6%). These results show that DOMIN expands the observed CATH homologous-superfamily co-occurrence space by proposing additional candidate pairings beyond those captured by TED-derived multidomain architectures. We then examined the distribution of these co-occurrences across CATH class combinations (Fig. 2d). Natural co-occurrences were unevenly distributed, with class combinations such as 1-3, 2-2, 2-3, and 3-3 contributing many pairs. DOMIN-predicted candidate novel co-occurrences showed a broadly similar class-level distribution, suggesting that they follow coarse-grained structural patterns present in natural multidomain architectures rather than being uniformly distributed across CATH classes. Finally, we visualized the CATH homologous-superfamily co-occurrence space as an adjacency matrix, where entries represent TED-derived natural co-occurrences and DOMIN-predicted candidate novel co-occurrences after removing pairs already observed in the TED-derived set (Fig. 2e). The inset network highlights local connectivity patterns among selected CATH homologous superfamilies. This representation illustrates how DOMIN expands the observed natural co-occurrence space with additional candidate pairings, which should be interpreted as hypotheses for potentially compatible or under-sampled domain combinations rather than confirmed natural co-occurrences.

We further evaluated the sequence identity and structural similarity (TM-score) between DOMIN-predicted paired domains and the corresponding ground-truth partners (see Section C.2). As shown in Fig. 2f, most predictions showed low sequence identity to the ground truth, whereas TM-scores separated into two main regimes. High-TM-score predictions indicate recovery of structurally similar partner domains despite limited sequence similarity, while low-TM-score predictions differ substantially from the annotated partners, suggesting that DOMIN can retrieve alternative candidate partners beyond close structural analogs of the ground truth. Fig. 2g shows a representative example. The query domain is shown as a blue backbone, and the ground-truth and predicted partners are shown in distinct colors. The TM-score matrix shows that the predicted partners are structurally distinct from the ground-truth partner and, in several cases, from one another. This illustrates the structural diversity of DOMIN-predicted candidate partners for the same query domain at the CATH homologous-superfamily level.

## 4 Designing multidomain proteins with DOMO

We first examined how training-set refinement and model scaling affect DOMO generation quality. On the held-out TED benchmark (Section A.1), filtering the training set to retain proteins with pLDDT greater than 70 improved performance relative to training on the full dataset. On DOMIN-retrieved domain-pair specifications (Section A.2), scaling the model from 200M to 1.2B parameters produced further gains (Fig. 3c). These results suggest that DOMO benefits from higher-confidence protein sequences and increased model capacity, despite being trained only on sequence data. We therefore used the 1.2B-parameter model trained on the refined dataset for the following analyses unless otherwise stated.

**Fig. 3.**
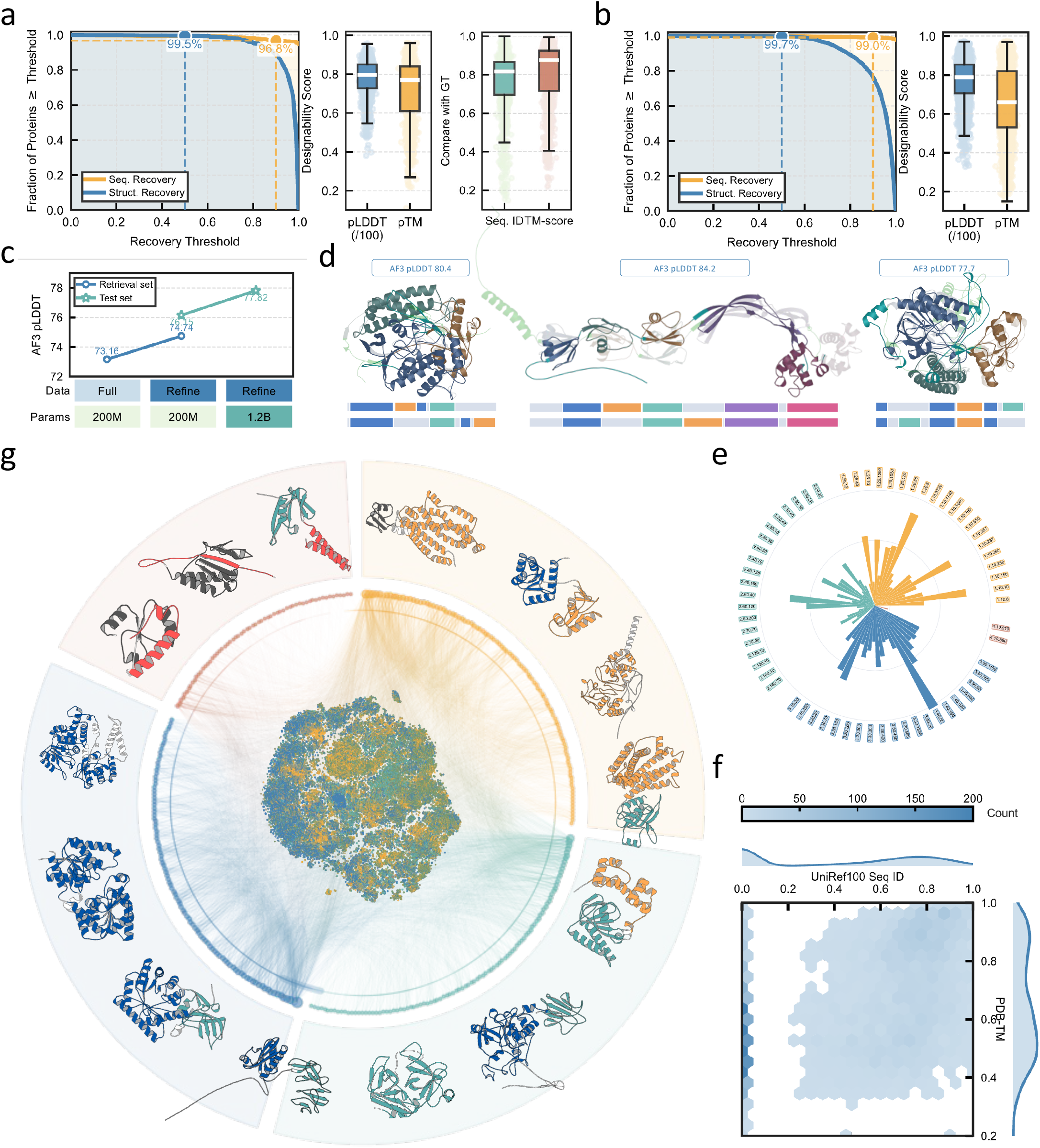
DOMO designs natural and novel multidomain proteins. **a**, Performance on the held-out TED test set, including sequence recovery, structure recovery, the distributions of pLDDT and pTM for designed proteins, and similarity to the native targets in sequence identity and TM-score. **b**, Quality assessment on a random sample of 5,000 proteins from the 5 million retrieval-derived designs. **c**, Effects of model scaling and data refinement on structural quality, measured by AlphaFold3 pLDDT on both the held-out test set and the DOMIN-retrieved set. The refined dataset retains only training proteins with pLDDT *>* 70. **d**, Representative examples of alternative domain organizations from the held-out TED test set. In these cases, DOMO realizes the target domains in an order different from the native protein. **e**, Counts of CATH labels in the 5 million retrieval-derived proteins. For clarity, only the most frequent CATH labels are shown: the top 20 for Classes 1–3 and the top 2 for Class 4. **f**, Novelty assessment of the 5,000 sampled retrieval-derived proteins. The joint distribution compares the highest sequence identity to known proteins in UniRef100 against the highest structural similarity (measured by TM-score) to PDB entries. **g**, Global view of the retrieval-derived design landscape, illustrating that the generated proteins organize into a structured multidomain recombination space featuring diverse representative architectures.

We next evaluated whether DOMO can realize domain-pair specifications as full-length multidomain proteins. On held-out TED multidomain proteins (Section A.1), DOMO was conditioned on annotated domain pairs, and the generated sequences were assessed for recovery of the input domains. For each generated protein, sequence and structural recovery were computed by matching generated domains to the input domains and averaging recovery across the input domains; the recovery curves therefore report the fraction of generated proteins whose averaged recovery exceeds each threshold (Section C.3). DOMO recovered the input domain pairs with high fidelity: 96.8% of the generated proteins achieved a sequence recovery of 0.9, and 98.1% achieved a structural recovery of 0.5 (Fig. 3a). The generated full-length proteins also showed high AlphaFold3-predicted confidence, with robust pLDDT and pTM distributions, and remained highly similar to the native targets in both sequence identity and TM-score.

Beyond reconstructing existing domain arrangements, DOMO also produced alternative organizations of the same domain set. In approximately 10% of the held-out TED test proteins, the generated sequences realized the target domains in an order different from that of the native protein (Fig. 3d). These designs nevertheless preserved the same constituent domains and formed plausible overall architectures, suggesting that DOMO is not limited to reproducing native domain orderings, but can also generate alternative organizations of the same domain set.

We then used DOMO to generate 5 million proteins conditioned on domain pairs produced by DOMIN. In a random sample of 5,000 proteins from this set, DOMO showed similar performance on DOMIN-retrieved pairs, with 99.0% of generated proteins achieving sequence recovery of 0.9 and 99.7% achieving structural recovery of 0.5 (Fig. 3b). The generated proteins also showed high pLDDT and pTM distributions, indicating that DOMO can convert DOMIN-retrieved domain-pair specifications into full-length sequences predicted to form confident multidomain structures.

We further assessed the novelty of proteins generated from DOMIN-retrieved domain pairs by comparing each generated protein with known proteins in sequence and structure space. For each generated protein, we computed its highest sequence identity to UniRef100 and its highest structural similarity to PDB structures, measured by TM-score (Section C.4). The generated proteins covered a wide range of similarities to known proteins (Fig. 3f), with many showing low sequence identity to UniRef100 while retaining moderate to high structural similarity to PDB entries. Together with the strong recovery and AlphaFold3-predicted confidence shown in Fig. 3b, these results suggest that DOMO can transform DOMIN-retrieved domain-pair specifications into plausible multidomain proteins that are not merely direct recapitulations of known sequences.

Finally, we examined the diversity and organization of the full set of 5 million proteins generated from DOMIN-retrieved domain pairs. The paired domains covered a broad set of CATH annotations (Fig. 3e), indicating broad coverage of domain classes among the generated combinations. At the global level, the generated proteins organized into a structured recombination landscape containing diverse representative multidomain architectures (Fig. 3g). Together, these results suggest that DOMINO can generate a large, diverse, and structured collection of plausible multidomain proteins from DOMIN-retrieved domain-pair specifications, enabling systematic exploration of multidomain architectures beyond known natural examples.

## 5 Discussion

Natural multidomain proteins are not arbitrary concatenations of structural modules, but exhibit recurring patterns of domain co-occurrence that likely reflect a mixture of evolutionary history, structural constraints, chain connectivity, and functional context. In this work, we used these patterns as a learnable prior for multidomain protein design. Rather than treating each domain independently or relying solely on manually specified domain architectures, DOMINO learns from observed domain associations and uses this information to guide the generation of full-length sequences containing specified domain combinations. The results show that domain co-occurrence provides useful information for proposing domain partners and for realizing these combinations at the sequence level, supporting the view that multidomain protein design can benefit from explicit modeling of architecture-level organization.

The proposed domain combinations generated by DOMINO should be interpreted as design hypotheses rather than validated biological entities. Natural co-occurrence provides evidence that certain domains have been observed together in the context of a single protein chain and may therefore represent compatible architectural units, but it does not define a complete set of allowable combinations. Similarly, the absence of a domain pair from current databases may reflect limited evolutionary sampling, annotation coverage, or dataset bias, rather than true incompatibility. By learning statistical regularities from observed multidomain proteins, DOMINO prioritizes candidate architectures that are informed by natural organization while still allowing exploration beyond directly observed co-occurrences. These candidates can then be evaluated by structural prediction, targeted computational analyses, and ultimately experimental characterization.

A major limitation of the present study is that all evaluations are in silico. Domain recovery, predicted structural confidence, structural similarity, and sequence novelty provide useful evidence that the generated proteins are computationally plausible, but they do not demonstrate that the proteins will express, fold, remain stable, or function in biological settings. AlphaFold-based confidence scores are useful for large-scale filtering and comparative evaluation, but they are not substitutes for experimental characterization. Future work should therefore include synthesis and biochemical testing of representative designs selected across different CATH classes, different degrees of sequence novelty, and different types of retrieved domain combinations. Measurements of expression, solubility, folding, thermal stability, oligomeric state, and, where appropriate, functional activity will be necessary to estimate the experimental success rate of the framework and to determine which computationally prioritized architectures are physically and biologically realizable.

A second important consideration is that multidomain protein design requires evaluation beyond global structure-confidence metrics. Whole-chain pLDDT, pTM, and global structural similarity are informative, but they can obscure the local features that determine whether a domain combination is architecturally coherent. For domain–domain combinations, the most critical regions often include domain boundaries, linkers, inter-domain interfaces, and the sequence context surrounding each domain. A design may have high average confidence while still containing an implausible linker, a strained domain boundary, or a poorly packed interface. Conversely, a protein with moderate global confidence may still recover the intended domains and place them in a plausible relative arrangement. Future evaluation should therefore emphasize multidomain-specific metrics, including domain-wise confidence, boundary quality, linker geometry, inter-domain contact consistency, interface confidence, and relative domain orientation. Such analyses will be essential for distinguishing proteins that merely contain recognizable domains from those that form coherent multidomain architectures.

The present work focuses its systematic evaluation on pairwise domain specifications, which provide a controlled setting for testing whether co-occurrence can be learned and converted into sequence-level designs. However, natural multidomain proteins frequently contain more than two domains, repeated modules, and higher-order architectural dependencies. The current framework may in principle be extended to such settings, but the behavior of the model on larger domain sets has not yet been systematically evaluated. Future studies should therefore examine whether learned pairwise compatibility patterns generalize to architectures with three or more domains, and whether additional conditioning information such as domain order, copy number, spacing, or desired inter-domain contacts improves the design of higher-order multidomain proteins. These extensions would allow a more complete assessment of multidomain organization beyond pairwise co-occurrence.

Finally, the large-scale designs generated in this study suggest a route for exploring multidomain architectures beyond those captured in natural protein databases. Natural databases record domain combinations sampled and retained by evolution, whereas DOMINO can propose additional combinations supported by learned co-occurrence patterns and favorable in silico properties. With further curation, such design sets could form the basis of future resources for multidomain design, providing prioritized candidates for studying domain recombination, benchmarking design methods, and selecting targets for experimental testing. Thus, generative design can complement natural domain databases by highlighting underexplored regions of multidomain protein space.

In summary, this work supports domain co-occurrence as a useful predictive prior for multidomain protein design. Although experimental validation and more detailed multidomain-specific evaluation remain necessary, the results demonstrate a scalable strategy for exploring new combinations of existing structural modules and generating full-length sequences that realize these candidate architectures.

## Supplementary Information

### A Data construction

#### A.1 Training, validation, and test datasets

**Fig. S1.**
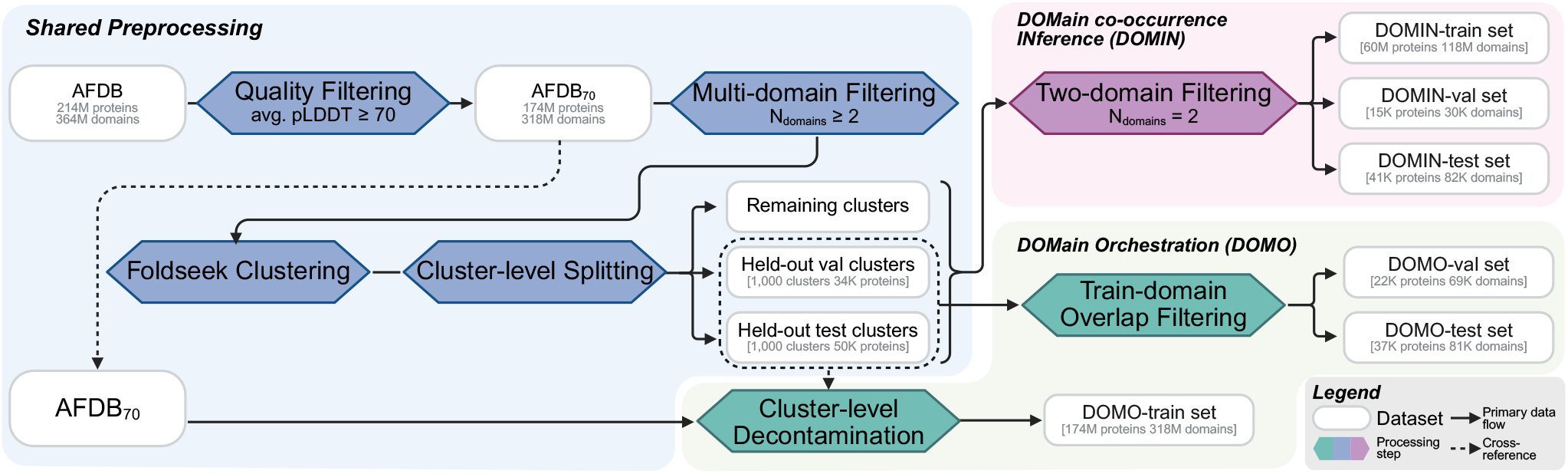
Overview of the data pipeline used to construct the data splits for DOMIN and DOMO.

We constructed the training data from TED (The Encyclopedia of Domains) [20], which provides systematic domain annotations for proteins in the AlphaFold Database (AFDB) [19]. As summarized in Fig. S1, the pipeline consists of a shared preprocessing stage followed by two task-specific branches, one for DOMIN learning and the other for DOMO.

In the shared preprocessing stage, we first removed proteins with mean pLDDT scores below 70 to reduce noise from low-confidence AlphaFold predictions, which can propagate to inaccurate TED domain segmentations. We then retained multidomain proteins and directly adopted the Foldseek clusters assignments [28]. Briefly, Foldseek clusters first grouped AFDB proteins by MMseqs2 sequence clustering, selected the highest-pLDDT representative from each cluster, and then constructed Foldseek clusters based on structural overlap. Clusters were split at the cluster level into training, validation, and test partitions, with 1,000 clusters reserved for validation and 1,000 clusters reserved for testing. This cluster-level split was used as the common starting point for both downstream branches.

For the DOMIN branch, we further restricted the data to proteins with exactly two annotated domains, matching the pairwise domain co-occurrence task used for retrieval. The resulting two-domain proteins were separated into training, validation, and test sets according to the cluster split defined above. Finally, to prevent information leakage, we removed from the validation and test sets any proteins containing a domain that also appeared in the training set.

For the DOMO branch, we instead retained the broader multidomain set defined by the shared preprocessing stage. The generation validation and test sets were taken from the held-out validation and test clusters described above. Similarly, to prevent information leakage, we removed from the validation and test sets any proteins containing a domain that also appeared in the generation training set.

The statistics of the dataset are summarized in Table S1.

#### A.2 Retrieval database

The retrieval database was constructed based on the training dataset construction (see Section A.1). Using Foldseek clusters described above, we obtained a protein database containing approximately 90 million proteins. We analyzed the distribution of proteins across clusters and observed a pronounced long-tail effect. To preserve this long-tail structure while obtaining a more compact retrieval database, we applied power-law sampling to proteins within each cluster with an exponent of 0.75, reducing the total protein count from 90 million to 16 million while effectively maintaining the clustering characteristics (see Fig. S2). The final retrieval pool comprised approximately 40 million domains.

**Table S1.** Statistics of the datasets used for DOMIN and DOMO.

| Split | DOMIN |  | DOMO |  |
| --- | --- | --- | --- | --- |
|  | Proteins | Unique Domains | Proteins | Unique Domains |
| Train | 60,090,343 | 117,768,291 | 173,837,555 | 318,041,112 |
| Validation | 15,432 | 30,395 | 22,173 | 69,461 |
| Test | 40,898 | 81,483 | 36,551<sup>a</sup> | 81,054<sup>b</sup> |

<sup>a</sup> Of the 36,551 proteins in the test split, 1,000 were used for the held-out design benchmark.
<sup>b</sup> The design benchmark subset spans 2,215 unique domains.

For the model-scaling and data-refinement analysis in Fig. 3c, we used a fixed development subset of 1,000 DOMIN-retrieved domain-pair specifications. This subset was used only for model selection and is distinct from the random sample of 5,000 proteins drawn from the final 5 million retrieval-derived designs in Fig. 3b.

#### A.3 Construction of the 5 million retrieval-derived domain pairs

To construct the 5 million retrieval-derived domain pairs used for large-scale generation, we started from the retrieval pool described above, which contains approximately 40 million domains. From this pool, we selected the subset of domains originating from proteins with exactly two annotated domains, yielding more than 20 million query domains.

For each query domain, DOMIN was then used to search the full 40 million-domain retrieval pool and retrieve the top 10 partner candidates. To construct the final set, we randomly selected 5 million query domains from this query set and, for each selected query, randomly chose one partner from its top 10 retrieved candidates. This yielded 5 million retrieval-derived domain pairs, with each query domain represented only once.

Each selected query–partner pair was then used as a conditioning specification for DOMO to generate one multidomain protein sequence. The random sample of 5,000 proteins analyzed in Fig. 3b and Fig. 3f was drawn from this final set of 5 million retrieval-derived designs.

### B Implementation details

#### B.1 DOMIN: domain co-occurrence inference

DOMIN learns a latent domain co-occurrence space in natural multidomain proteins. We adopted InfoNCE loss [29] for contrastive learning. The InfoNCE loss can be detailed as:

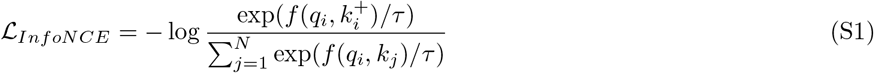

Here, *q*_*i*_ and 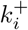 represent the query and positive key embeddings of the *i*-th domain pair in a batch, respectively. *f* (*q*_*i*_, *k*_*j*_) is the similarity score between two embeddings, which in this study is defined as the cosine similarity between *q*_*i*_ and *k*_*j*_. *N* is the total number of domain pairs in a batch and *τ* is a learnable temperature parameter initialized at 0.07. During training, samples within each batch were randomly selected. In each batch, the positive sample corresponds to the two domains that co-occur within the same protein, while all remaining domain pairs (i.e., (*q*_*i*_, *k*_*j*_) where *j*≠ *i*) are treated as negative samples. This constitutes a standard contrastive learning framework.

**Fig. S2.**
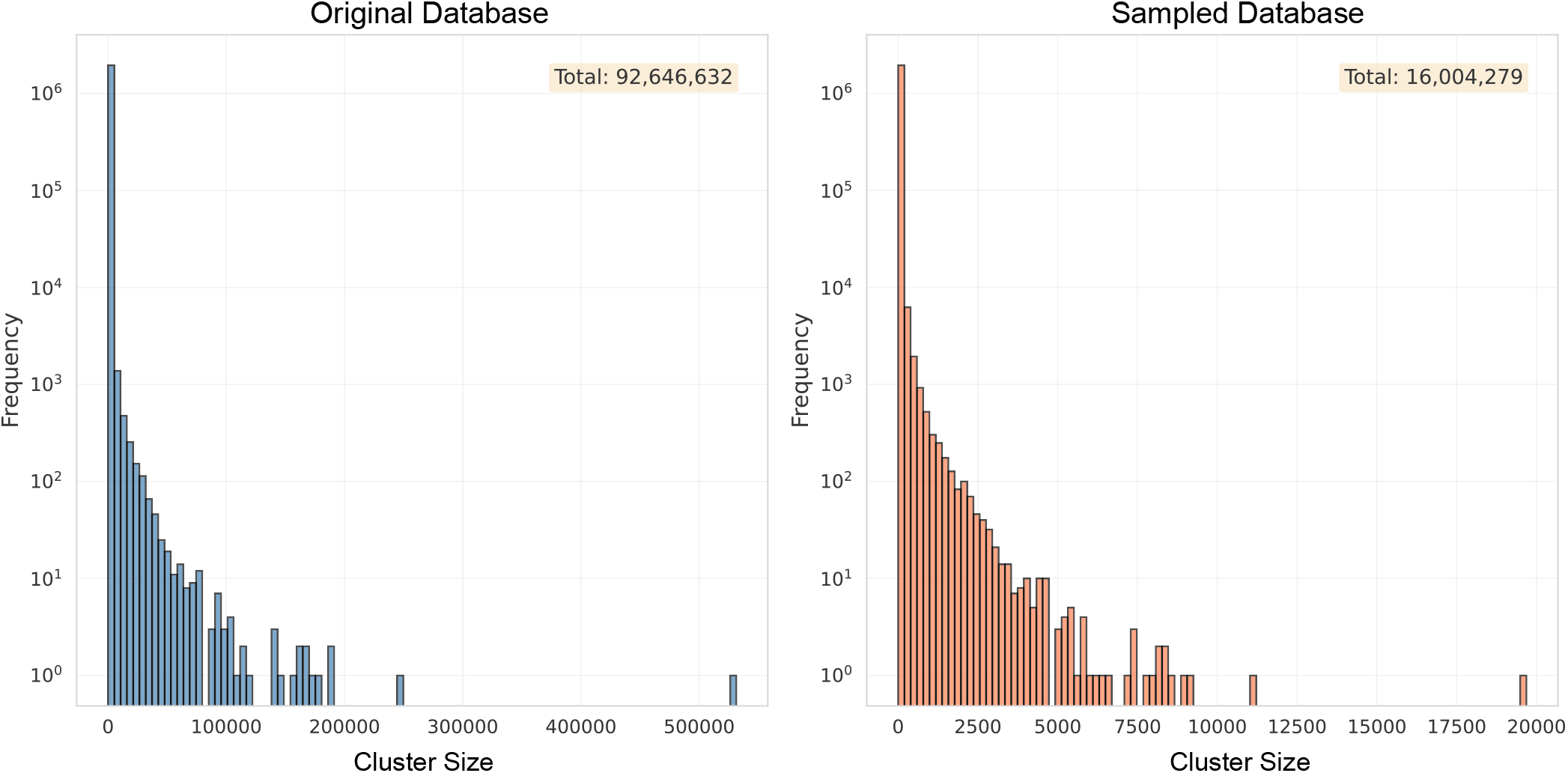
Protein count distribution per Foldseek clusters before and after power-law sampling.

We used the DeepSpeed strategy with ZeRO-2 optimization [30] and employed the AdamW optimizer [31], setting *β*_1_ = 0.9, *β*_2_ = 0.98 and we utilized *L*_2_ weight decay of 0.01. We gradually increased the learning rate from 0 to 1e-4 over the first 2000 steps and decreased it to 2e-5 using the cosine annealing schedule. The overall training phase lasted approximately 300K steps trained on 16 NVIDIA 80G H800 GPUs.

We truncated domain sequences to a maximum of 1024 tokens. When the length of a domain exceeded 1024 amino acids, we randomly selected a starting position and extracted a consecutive segment of 1024 amino acids as input for DOMIN. Our total batch size consisted of 1024 domain pairs. Additionally, we employed mixed-precision training [32] for DOMIN.

#### B.2 DOMO: domain orchestration

For DOMO, we used a Transformer-based autoregressive model [33], in which ESM2 [34] serves as the domain feature encoder. The decoder adopts a modern dense design [35], including grouped-query attention [36] with QK-Norm [37], SwiGLU feed-forward layers [38], rotary positional embeddings [39], and RMSNorm [40] with pre-normalization, while being augmented with cross-attention to domain-derived conditioning features.

Each input domain sequence was first encoded independently by the domain feature encoder to obtain contextual residue-level embeddings. For each protein, the embeddings of all input domains were reassembled at the protein level, concatenated after removing intra-domain padding tokens, and then truncated or padded to a fixed length. The resulting embeddings were projected by a learned linear layer from the encoder hidden dimension to the decoder hidden dimension, and together with its attention mask were supplied to the autoregressive decoder as cross-attention memory. This design enables DOMO to attend to residue-level features from multiple domains while generating the full-length multidomain protein sequence.

The model was trained under a standard autoregressive language-modeling objective:

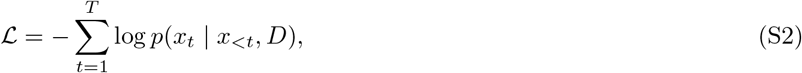

where *x*_*t*_ is the target residue at position *t, x*_*<t*_ denotes the preceding residues, and *D* = {*d*_1_, *d*_2_, …, *d*_*N*_ } denotes the set of input protein domains.

We trained DOMO using DeepSpeed ZeRO-2 optimization [30], Flashattention-2 [41], and mixed-precision training [32]. Optimization was performed with AdamW [31] using *β*_1_ = 0.9, *β*_2_ = 0.98, and a weight decay of 0.01. The learning rate was linearly warmed up to 1 × 10^−4^ during the first 1,000 steps and then cosine-decayed to 1 × 10^−5^. Training was run for 200K steps on 8 NVIDIA 80G H800 GPUs with a global batch size of 512 and a maximum sequence length of 1024 tokens.

### C In silico evaluation

#### C.1 Retrieved domain pairs are more designable than random pairs

To assess whether retrieval improves the designability of domain combinations, we compared DOMIN-retrieved domain pairs with several random-pairing baselines.

##### Pure random sampling

For each query domain, we randomly sampled a partner domain from the full retrieval pool (Section A.2), without imposing any constraint on structural similarity or natural co-occurrence.

##### CATH-level stratified random sampling

To examine whether structural similarity alone can account for designability, we constructed random baselines at four levels of the CATH hierarchy. For each query domain, we identified its native partner domain and then randomly sampled a domain matching that partner at one of the following levels:

- **Class (C):** same Class (e.g., C1)
- **Architecture (A):** same Architecture (e.g., C1.A1)
- **Topology (T):** same Topology (e.g., C1.A1.T1)
- **Homology (H):** same Homology superfamily (e.g., C1.A1.T1.H720)

Among the 5,000 query domains, 3,452 had native partners with CATH annotations and were therefore eligible for CATH-level stratified sampling. For these queries, the mean candidate-pool sizes were approximately 11 million, 4 million, and 1 million domains at the Class, Architecture, and Topology levels, respectively. For the Homology-level baseline, 2,585 of the 3,452 queries had a non-empty H-level candidate pool, with a mean pool size of approximately 160 thousands domains per query. For the remaining 867 queries, no H-level pool was available and we therefore reverted to Topology-level sampling. Queries whose native partners lacked CATH annotations were instead assigned partners by pure random sampling from the retrieval pool.

Sampling was performed independently for each query domain, yielding 5,000 query–partner pairs for each strategy. Using the same set of query domains, the DOMIN baseline was constructed by taking the top-ranked partner retrieved by DOMIN for each query. DOMO was then used to generate full-length proteins from each query–partner specification, and the resulting designs were evaluated using AlphaFold3 (AF3) pLDDT and pTM.

**Fig. S3.**
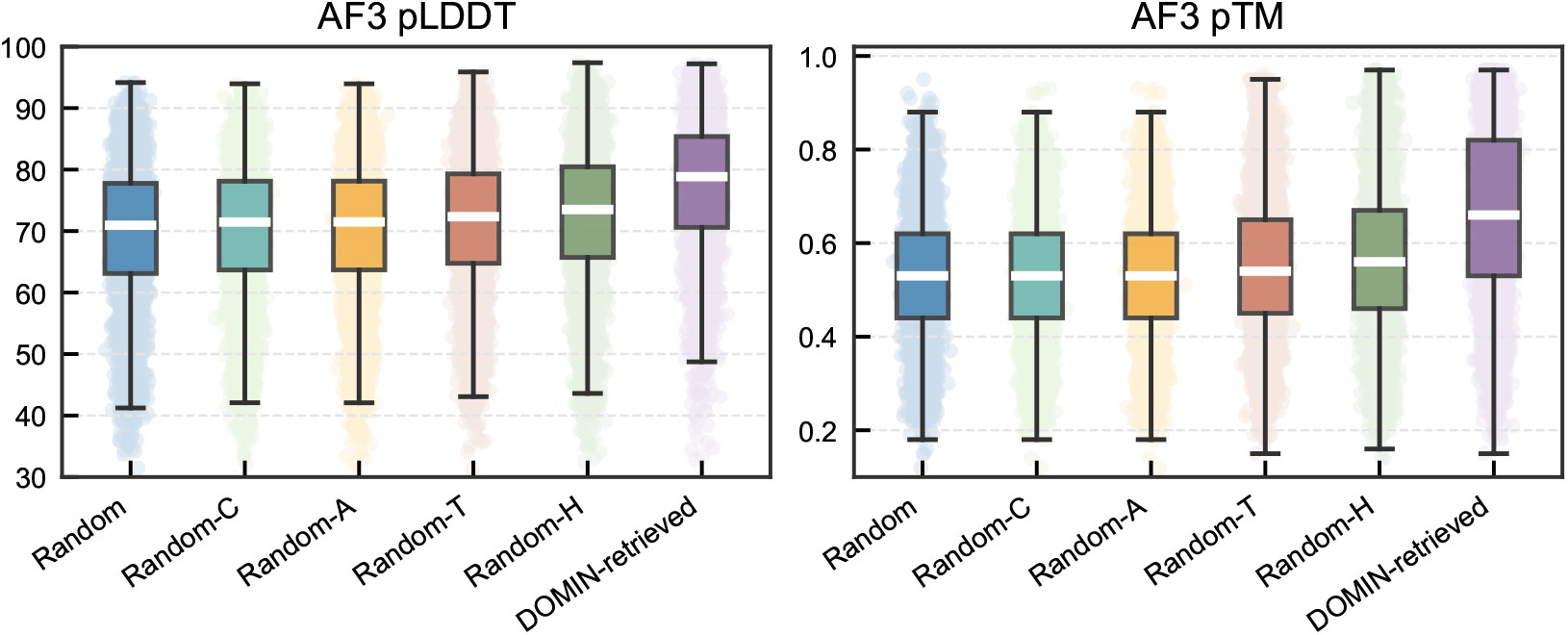
Retrieval-derived domain pairs are more designable than randomly paired domains. Distributions of AF3 pLDDT and AF3 pTM for proteins generated from pure random pairs, CATH-level stratified random pairs, and DOMIN-retrieved pairs. Random-C, Random-A, Random-T, and Random-H denote random partners sampled at the Class, Architecture, Topology, and Homology levels, respectively.

Across both metrics, proteins generated from DOMIN-retrieved pairs showed higher predicted structural confidence than those generated from randomly paired domains (Fig. S3). Increasing the CATH specificity of the random baselines led to only marginal improvement, indicating that structural similarity to the native partner alone is insufficient to explain the improved designability. These results suggest that DOMIN captures compatibility signals that help DOMO generate coherent multidomain proteins.

#### C.2 Cluster-based evaluation of predicted domain pairs

To evaluate the predicted domain co-occurrences, we clustered domains in the retrieval database at 90% sequence identity, where each cluster contains multiple domains with different ground truth paired domains. Each cluster representative was used as a query domain, and the top-1 retrieved domain was compared against all corresponding ground truth paired domains within the same cluster. The maximum sequence identity and maximum TM-score were retained for analysis.

#### C.3 Domain recovery metrics

To quantify how well a designed protein realizes its target domains, we defined domain recovery metrics in both sequence and structure space. Because some TED domains are discontinuous in the primary sequence, a target domain may correspond to one or more sequence intervals even though these intervals jointly form a single structural domain. The recovery metrics below therefore operate at the domain level rather than treating each target domain as a single contiguous sequence segment.

For sequence-based domain recovery, each target domain was represented as the union of its constituent sequence segments. Each segment was matched against the designed full-length protein sequence using local pairwise alignment, and the number of identical residues was accumulated across all segments belonging to the same domain. The sequence recovery score for a domain was then defined as the total number of matched residues divided by the total domain length. Protein-level sequence recovery was computed by averaging this score across all target domains in the same design.

For structure-based domain recovery, each target domain was first mapped back to its corresponding residue interval or intervals in the reference protein sequence, and the associated structural fragments were extracted from the reference structure. When a domain comprised multiple sequence intervals, these fragments were combined to form a single reference domain structure. We then aligned this reference domain structure against the designed full-length predicted structure using TMalign.

For each domain, the structural recovery score was defined as the TM-score normalized by the target domain length rather than by the full protein length. Protein-level structural recovery was then computed by averaging these domain-level TM-scores across all target domains in the same design.

Unless otherwise stated, the sequence and structure recovery values reported in the main text correspond to these protein-level averages, further aggregated across all proteins in the evaluated set. Both metrics were implemented in custom analysis scripts.

##### Algorithm 1

Sequence-based domain recovery

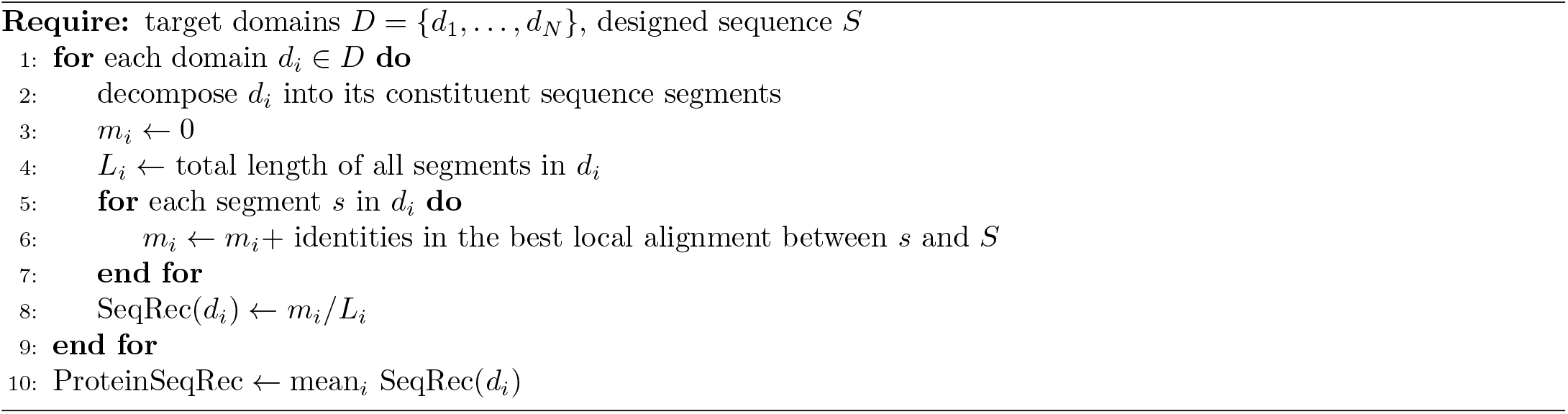

##### Algorithm 2

Structure-based domain recovery

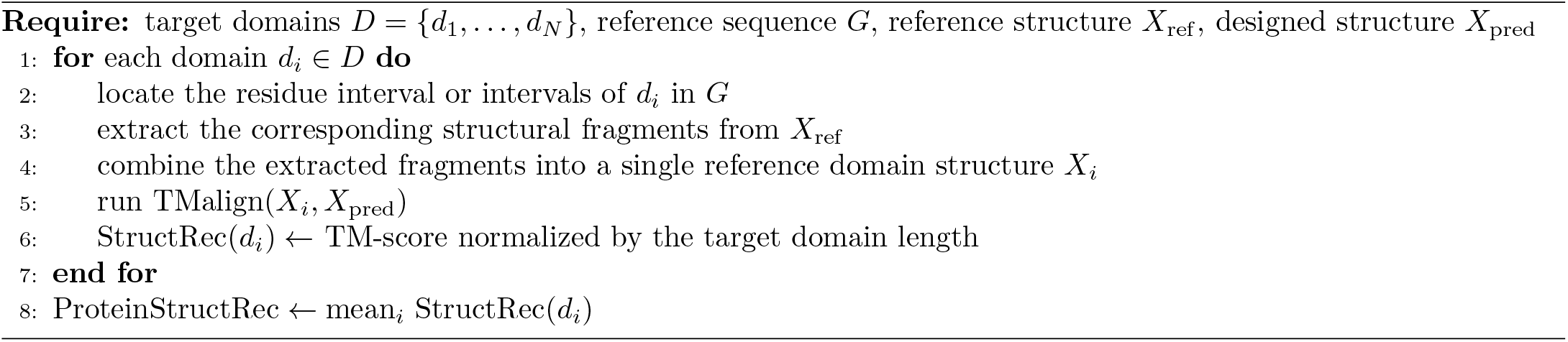

#### C.4 Protein novelty assessment

Protein novelty was assessed in both sequence and structure space. For sequence novelty, each designed protein was searched against UniRef100 [42] using MMseqs2 [43], and novelty was quantified relative to the closest sequence matches in the database. For structure novelty, each designed protein structure was searched against the Protein Data Bank (PDB) [16] using Foldseek, and novelty was quantified relative to the closest structural matches returned by the search.

For protein novelty assessment, we compared each designed protein against known natural proteins in both sequence and structure space. Sequence-based novelty was evaluated by searching the designed sequences against UniRef100 with MMseqs2 and using the closest sequence matches to quantify similarity in sequence space. Structure-based novelty was evaluated by searching the predicted structures of the designed proteins against the PDB with Foldseek and using the closest structural matches to quantify similarity in structure space. In the main text, UniRef100 sequence identity and PDB-TM refer to these nearest-neighbor similarities in sequence and structure space, respectively. We considered designs with UniRef100 sequence identity below 0.3 and PDB-TM below 0.5 to be novel.

The sequence- and structure-based novelty searches were implemented using the following command lines.

##### Software versions

~~~
MMseqs2 1668032cda9c0e1c5e1aab559aa665ee6973c7e6;Foldseek 10.941cd33.
mmseqs createdb query_fasta_path query_db
mmseqs search query_db ur100_db_path result_db tmp_dir \
--gpu 1 --min-seq-id 0 --alignment-mode 3 --max-seqs 100 \
-s 1 -c 0.8 --cov-mode 0 --threads 120 -a
mmseqs align query_db ur100_db_path result_db alnNew -a --threads 120
mmseqs convertalis query_db ur100_db_path alnNew align_fasta_path \
--format-output query,target,qseq,tseq,pident,fident,nident \
--threads 120
~~~

~~~
foldseek easy-search query_dir pdb output.txt tmpFolder \
--alignment-type 1 \
--format-output query,target,qtmscore,ttmscore,alntmscore,lddt \
--tmscore-threshold 0.0 \
--exhaustive-search \
--max-seqs 10000000000 \
--threads 120
~~~

## Data availability

The pre-trained DOMINO weights can be downloaded from https://huggingface.co/westlake-repl/DOMINO.

## Code availability

DOMINO is open-sourced under the MIT license. The code repository is available at https://github.com/westlake-repl/DOMINO. The DOMINO web server is located at http://www.protein-domino.com/.

